# Serosurveillance and the first detection of Bourbon virus RNA in a wildlife host

**DOI:** 10.1101/2024.06.04.597417

**Authors:** Gayan Bamunuarachchi, Fernando Najera, Ishmael D. Aziati, Jamie L. Palmer, Elizabeth G. Biro, Dave Wang, Sharon L. Deem, Adrianus C. M. Boon, Solny A. Adalsteinsson

## Abstract

Bourbon virus (BRBV) is an emerging pathogen that can cause severe and fatal disease in humans. BRBV is vectored by *Amblyomma americanum* (lone star ticks), which are widely distributed across the central, southern, and eastern United States. Wildlife species are potentially important for the maintenance and transmission of BRBV, but little is known about which species are involved, and what other factors play a role in the exposure to BRBV. To assess the exposure risk to BRBV among wildlife in the St. Louis area, we collected sera from 98 individuals, representing 6 different mammalian species from two locations in St. Louis County: Tyson Research Center (TRC) and WildCare Park (WCP) from fall 2021 to spring 2023. The sera were used in a BRBV neutralization assay to detect neutralizing antibodies and RT-qPCR for viral RNA analysis. We also sampled and compared the abundance of *A. americanum* ticks at the two locations and modeled which factors influenced BRBV seropositivity across species. In TRC, we observed a high rate of seropositivity in raccoons (*Procyon lotor*, 23/25), and white-tailed deer (*Odocoileus virginianus*, 18/27), but a low rate in opossums (*Didelphis virginiana*, 1/18). Neutralizing antibodies were also detected in sampled TRC bobcats (*Lynx rufus*, 4/4), coyotes (*Canis latrans*, 3/3), and a red fox (*Vulpes vulpes*, 1/1). The virological analysis identified BRBV RNA in one of the coyote serum samples. In contrast to TRC, all sera screened from WCP were negative for BRBV-specific neutralizing antibodies, and significantly fewer ticks were collected at WCP (31) compared to TRC (2,316). Collectively, these findings suggest that BRBV circulates in multiple wildlife species in the St. Louis area and that tick density and host community composition may be important factors in BRBV ecology.

## INTRODUCTION

Emerging and reemerging infectious diseases account for one-third of global human mortalities, and viruses cause a significant number of these infectious diseases. Many of these viruses are vectored via arthropods. Ticks are the number one disease-causing vector in the United States (US) [1-3] and they vector viral pathogens including Powassan virus, Colorado tick fever virus, Heartland virus (HRTV), and Bourbon virus (BRBV). BRBV belongs to the family *Orthomyxoviridae*, genus *Thogotovirus*. This novel virus was first isolated in 2014 from a fatal human infection following a tick bite in Bourbon County, Kansas, US [4]. To date, five BRBV human infections have been reported from Kansas, Oklahoma, and Missouri [5]. While reported human BRBV cases are clustered in the central US, suspected cases have emerged elsewhere such as in New York, where a tick with BRBV RNA was removed from a resident who recovered from BRBV-like symptoms [6]. A human serological study in St. Louis, Missouri showed that 0.3-0.7% of individuals are seropositive for neutralizing antibodies against BRBV [7], suggesting that human infections with BRBV are more common and geographically widespread than previously recognized.

BRBV has been primarily isolated from lone star ticks (*Amblyomma americanum)* [6, 8-11], an aggressive human and animal biting tick, which is abundant and widely distributed across the central, southern, and eastern US [12, 13]. Several tick surveillance studies after the first BRBV human case found BRBV in lone star ticks in Kansas, northwestern Missouri, and eastern-central Missouri [9, 11, 14]. Since then, BRBV has been found in lone star ticks outside of the Midwest US, in New York, New Jersey, and Virginia [6, 10, 15]. Furthermore, Cumbie *et al* 2022 detected BRBV in a nonnative, invasive tick species, the long-horned tick (*Haemaphysalis longicornis*), that has been present in the US since at least 2014 [15].

How BRBV is maintained in nature remains unknown, but serosurveillance studies have identified some mammalian species that are permissive to BRBV infection. *Amblyomma americanum* ticks are generalist, 3-host ticks that readily feed on a wide range of vertebrate host species [16, 17]; therefore, many species are potentially exposed to BRBV or involved in its enzootic transmission. In northwestern Missouri, BRBV-neutralizing antibodies were detected in serum samples from domestic dogs (*Canis lupus familiaris)*, Eastern cottontails (*Sylvilagus floridanus)*, horses (*Equus caballus)*, raccoons (*Procyon lotor)*, and white-tailed deer (*Odocoileus virginianus*) [18]. BRBV-neutralizing antibodies were also found in white-tailed deer serum collected from North Carolina and New York [6, 19]. While these studies are critical first steps to understanding BRBV ecology, many used banked serum samples collected across wide geographic ranges where information is lacking about tick density and BRBV prevalence in local tick populations; therefore, negative results are ambiguous in that they could represent an absence of ticks, BRBV, or host resistance to BRBV infection. To understand how BRBV is transmitted within wildlife host communities, and which species are permissive, it is important to sample in areas with known BRBV prevalence. Furthermore, little is known about how BRBV seropositivity varies with ecological factors within a community such as host species, host age, host sex, and tick density. Finally, many wildlife species have not been tested for BRBV antibodies and viral RNA, limiting our overall understanding of BRBV ecology.

Here, we collected sera from six different mammal species and sampled free-living ticks at two different locations in St. Louis County, Missouri, and tested for the presence of BRBV-neutralizing antibodies and viral RNA. BRBV antibodies were detected in multiple different species including red foxes, bobcats, and coyotes. Furthermore, viral RNA was detected in one coyote serum sample. We identified tick density, wildlife species, and year of sampling as factors associated with the risk of exposure to BRBV.

## MATERIALS AND METHODS

### Study area

Wildlife serum samples and ticks were collected from two locations in St. Louis County, Missouri: Washington University’s Tyson Research Center (TRC, 38°31ʹN, 90°33ʹW) and the Saint Louis Zoo’s WildCare Park (WCP, 38°48ʹN, 90°11ʹW). TRC is an 800-ha environmental field station at the northeastern edge of the Ozarks ecoregion. Approximately 85% of TRC is forested, consisting of mainly deciduous oak-hickory forests on steep slopes and ridges. South-facing slopes are dominated by chinquapin oak (*Quercus muehlenbergii*) and eastern red cedar (*Juniperus virginiana*); protected slopes by flowering dogwood (*Cornus florida)*, white oak (*Q. alba*), and black oak (*Q. velutina)*; and bottomlands by slippery elm (*Ulmus rubra*) and American sycamore (*Plantanus occidentalis*) [20]. The forest understory contains areas of woody shrubs including spicebush (*Lindera benzoin)*, common buckthorn (*Frangula caroliniana*), and pawpaw (*Asimina triloba*). Throughout TRC there are large and patchy areas of Amur honeysuckle (*Lonicera maackii)* and *Sericea lespedeza* invasion, some of which are under ongoing management. TRC maintains an old field habitat in its central valley and has ∼24 ha of limestone/dolomite glades that are heavily invaded by red cedar. *Amblyomma americanum* is the most commonly encountered tick species at TRC; *Dermacentor variabilis* and *Ixodes scapularis* are also present [21]. Both Bourbon and Heartland viruses have been detected in field-collected *A. americanum* from TRC in multiple years [11].

WildCare Park (WCP) is a 172-ha, fenced property that was under active development at the time of the study for conversion from a country club-style property to a Saint Louis Zoo safari park. The property exemplifies a mosaic of bottomland habitats. It includes open grassland (part of an old golf course) and bottomland hardwood. Most sides of the property are surrounded by the Spanish Lake community. The east side of the property borders the Columbia Bottom Conservation Area, a 17.5 km^2^ floodplain located at the confluence of the Mississippi and Missouri rivers, which includes shallow wetlands, bottomland hardwoods, prairie, and cropland. Anecdotally, researchers at WCP report low encounter rates with ticks but prior to this study there have not been any systematic surveys to document tick density or abundance.

### Serum collections from wildlife species

During two winter field seasons (October 2021 – March 2022, and October 2022 – March 2023), we trapped medium to large mammals at TRC and WCP using cage traps and cable restraints. The protocol and procedures employed for the capture of wildlife species were approved by the Saint Louis Zoo Institutional Animal Care and Use Committee protocol #21-15. All species but coyotes were captured using commercial cage traps (Tomahawk models 108 and 207, Tomahawk Live Trap Co., Tomahawk, Hazelhurst, WI, USA), camtrip cage traps (Camtrip Cages, Bartsow, California, USA), and hand-made double guillotine door cage-traps, baited with road-killed deer, cottontail, squirrel, and/or a mixture of visual attractants and lures. Coyotes were captured using cable restraint devices (Freemont snare, Snareshop Co. Lidderdale, Iowa, USA). All animals were anesthetized with a combination of medetomidine-tiletamine-zolazepam and partially reversed with atipamezole (Nájera et al., in preparation). Up to 10 mL of blood was extracted by jugular (raccoons, Virginia opossums), cephalic (coyotes, bobcats, and red foxes), and ventral caudal (Virginia opossums) venipuncture. Blood was collected in EDTA-coated tubes and serum separator tubes. Blood collected in serum separator tubes was allowed to clot and was then centrifuged at 3,000 rpm for 10 min. The serum was removed and stored at -80°C until analysis.

Deer hunters at TRC provided blood samples and ticks from deer that they harvested under their individual licenses and tags from the Missouri Department of Conservation. In advance of the hunting season, we provided each hunter with sample collection kits that included instructions, gloves, disposable pipettes, and serum tubes for blood sample collection, and fine-tipped forceps and vials for tick collection. Upon collection, samples were transported to a refrigerator (blood) and -80°C freezer (ticks) for storage.

### Virus and cell culture

Vero E6 cells (ATCC) were grown in DMEM media (Corning Cellgro) supplemented with 10% FBS (Biowest), 2 mM L-Glutamine (Corning), 100 U/mL Penicillin (Life Technologies), 100 μg/mL streptomycin (Life Technologies), and 1x MEM vitamins (Corning). BRBV (strain BRBV-STL) [5] was grown on Vero E6 cells. Virus stocks were titrated in Vero E6 cells and sequenced and verified by the next generation. Subsequently, the virus was aliquoted, stored at -80°C, and used for all subsequent studies.

### BRBV neutralization assay

A focus reduction neutralization assay (FRNT) was developed for BRBV [7]. Briefly, Vero E6 cells were seeded overnight in 96-well plates (Corning). Heat-inactivated wildlife sera were diluted 1:20 in serum-free DMEM (DMEM-SF) and subsequently serially diluted 3-fold to a final dilution of 1:14580. Next, an equal volume of DMEM-SF containing 200 focus forming units (FFU) of BRBV-STL was added to each serum dilution and incubated for 1 h at RT. The addition of the virus resulted in a final serum dilution of 1:40 to 1: 29160. Next, the cells were washed once with PBS and the antibody-virus complexes were transferred onto cells and incubated for 1 h at 37°C.

Each serum sample and dilution was tested in duplicate and each assay included positive (pooled convalescent sera from BRBV-infected mice), and no serum controls. After 1 h at 37°C, the cells were washed once with PBS, and overlay media (100 µL of MEM containing 2% FBS and 1% methylcellulose) was added to each well. After 48 h at 37°C, the cells were fixed by adding 100 µL of 10% formalin on top of the overlay (final concentration 5% formalin) for 30 min at RT. Subsequently, cells were washed with PBS and the assay was developed and analyzed as described previously [7].

### RNA Extraction and Virus Detection in sera

RNA extraction was done by adding 20 µL of inactivated sera to the lysis buffer mix and beads from the MagMax viral pathogen nucleic acid isolation kit (Cat. # A48310, Thermo Fisher) following the manufacturer’s protocol. The mixture was then loaded onto the KingFisher Flex RNA extraction platform (Thermo Fisher Scientific). The extracted RNA was stored at -80°C until ready to be used. All RNA samples were screened in a one-step RT-qPCR, for the presence of BRBV viral RNA with two different primer/probe sets targeting BRBV NP and PB1. A sample was scored as positive if it had a Ct value equal to or less than 35 (Ct ≤ 35) with both primer/probe sets as previously reported [11]. Segment-specific RT-PCR was performed to amplify whole gene segments for sequencing by first synthesizing cDNA using SuperScript III (Invitrogen) according to the manufacturer’s instructions. The first-strand cDNA was then amplified in a PCR reaction with Phusion High Fidelity PCR Master Mix (New England Biolabs) using BRBV segment-specific primers targeting the segment termini. PCR amplicons were separated on 1% agarose gel and products of the correct size were gel excised and column purified for sanger sequencing using the respective segment-specific primers.

### Tick sampling

To compare *A. americanum* population size between the two sites, we sampled ticks with CO_2_-baited traps. CO_2_-baited traps are a preferred method for capturing *A. americanum* because they take advantage of the tick’s aggressive host-seeking behavior and they are unbiased by differences in habitat type/structure [20, 22]. We set up trapping arrays of 5 traps, placed 20m apart, at each of 4 locations at both TRC and WCP. We baited and collected from each trap twice during June 2023, for a total of 80 trap-days across TRC and WCP. Each trap consisted of a white textured cloth held down to the ground by 4 galvanized washers with ½” interior diameter placed at each corner. We put 300g of dry ice in a 16-oz plastic Tupperware container with holes drilled on the sides and placed the container upside down at the center of the white cloth. After two hours, we returned to collect the cloth by folding it into a 2-gallon zip lock bag, taking care to include any extra ticks that might be found on the washers or the plastic container. Zip lock bags with live ticks were transported to a -80C freezer for storage. All collected ticks were sorted and identified on cold tables to species and life stages using microscopy [23-26].

### Statistical analyses

BRBV neutralizing activities were analyzed by using GraphPad Prism version 10.2 software. IC_50_ values were calculated by using log (inhibitor) vs. response in variable slope (four parameters) fixing both bottom and top constrain at 0 and 100 respectively. IC_50_ value of more than 1:40 is considered positive. To understand the drivers of BRBV seropositivity among our wildlife sample cohort, we used generalized linear models with a logit link function and AIC model selection [27] in R Statistical Software (v4.3.2, [28]). We modeled the influence of four predictor variables – wildlife species, trapping season, age of the animal, and sex of the animal – on the odds that an animal tested positive for BRBV antibodies, as defined by the threshold above. Our response variable was a binary 0 or 1 to indicate a BRBV-negative or positive result, respectively. We limited the analysis to three species from TRC, for which we had sample sizes >5 individuals: Virginia opossum *(Didelphis virginiana)*, raccoon, and white-tailed deer *(Odocoileus virginianus*). We focused on TRC because of its relatively larger tick population, the known presence of BRBV circulating among ticks [11], and the greater sample size of wildlife hosts. We constructed a set of 11 candidate models that included null, global, and models with various combinations of the predictor variables that we expected could be important drivers of BRBV seropositivity (**Table 4**). We compared models with Akaike’s Information Criterion (AICc), which balances over vs. underfitting to determine the best candidate model to explain the data, corrected for small sample sizes, using package AICcmodavg [29]. We examined residual plots to assess model fit and generated predictions from the best-fit model(s) to understand the influence of predictor variables.

## RESULTS

### Wildlife capture results

In season 1 (Fall 2021-Spring 2022), we collected 46 serum samples in total from: raccoons (n = 12), Virginia opossums (n = 17), bobcats (*Lynx rufus*, n = 2), red foxes (*Vulpes vulpes*, n = 2), and white-tailed deer (n = 13) from TRC and WCP (Table 1). In season 2 (Fall 2022- Spring 2023), we captured and collected 52 serum samples from: raccoons (n = 21), Virginia opossums (n = 12), bobcats (n = 2), coyotes (*Canis latrans*, n = 3), and white-tailed deer (n = 14) from the same locations (Table 1). Most of the sampled animals were adults (77%), and males (62%). Only six sera were collected from juveniles and 17 from sub-adult animals (Table 1). None of the red foxes, coyotes, bobcats, and white-tailed deer were trapped in both seasons.

**Table 1:** Number, sex, and age of animal species tested for BRBV antibodies and viral RNA.

| Season | Location | Common name | # of animals captured | Sex composition (M/F) | Age composition (juvenile/sub adult/adult) |
| --- | --- | --- | --- | --- | --- |
| 1 (fall 2021 - spring 2022) | TRC | Virginia opossums | 7 | 4/3 | 0/1/6 |
|  |  | Red fox | 1 | 1/0 | 0/1/0 |
|  |  | Bobcat | 2 | 2/0 | 0/0/2 |
|  |  | Raccoon | 10 | 8/2 | 0/0/10 |
|  |  | White-tailed deer | 13 | 6/7 | 2/4/7 |
|  | WCP | Virginia opossums | 10 | 3/7 | 0/1/9 |
|  |  | Red fox | 1 | 1/0 | 0/1/0 |
|  |  | Raccoon | 2 | 2/0 | 0/0/2 |
| 2 (fall 2022 - spring 2023) | TRC | Virginia opossums | 11 | 8/3 | 0/2/9 |
|  |  | Bobcat | 2 | 0/2 | 1/1/0 |
|  |  | Coyote | 3 | 0/3 | 0/1/2 |
|  |  | Raccoon | 15 | 13/2 | 0/3/12 |
|  |  | White-tailed deer | 14 | 7/7 | 3/1/10 |
|  | WCP | Virginia opossums | 1 | 1/0 | 0/1/0 |
|  |  | Raccoon | 6 | 5/1 | 0/0/6 |

### Neutralization of BRBV by wildlife serum samples

Wildlife sera collected at TRC and WCP were subjected to virus-neutralizing assays to detect BRBV-neutralizing antibodies in the serum. In season 1, BRBV-neutralizing antibodies were detected in 1/1 red fox, 2/2 bobcats, 9/10 raccoons, and 7/13 white-tailed deer in TRC (Table 2). Importantly, only 1/7 Virginia opossums show marginally positive BRBV-neutralizing antibodies. In season 2, BRBV-neutralizing antibodies were detected in 2/2 bobcats, 3/3 coyotes, 14/15 raccoons, and 11/14 white-tailed deer captured in TRC (Table 2). All 11 opossum serum samples were negative for BRBV-neutralizing antibodies. Jointly in seasons 1 and 2, BRBV neutralizing antibodies were detected in 1/1 red fox, 4/4 bobcat, 3/3 coyote, 23/25 raccoon, 18/27 white-tailed deer, and 1/18 Virginia opossum serum samples in TRC (Figure 1A). The IC_50_ for the positive sera ranged between 1:100 and 1:6000 (Figure 1A). While we found high rates of BRBV seropositivity in wild animal serum collected at TRC (Figure 1A), serum samples collected from red fox (0/1), raccoons (0/8), and Virginia opossums (0/11) in WCP in both seasons were negative for BRBV-neutralizing antibodies (Figure 1B).

**Table 2:**
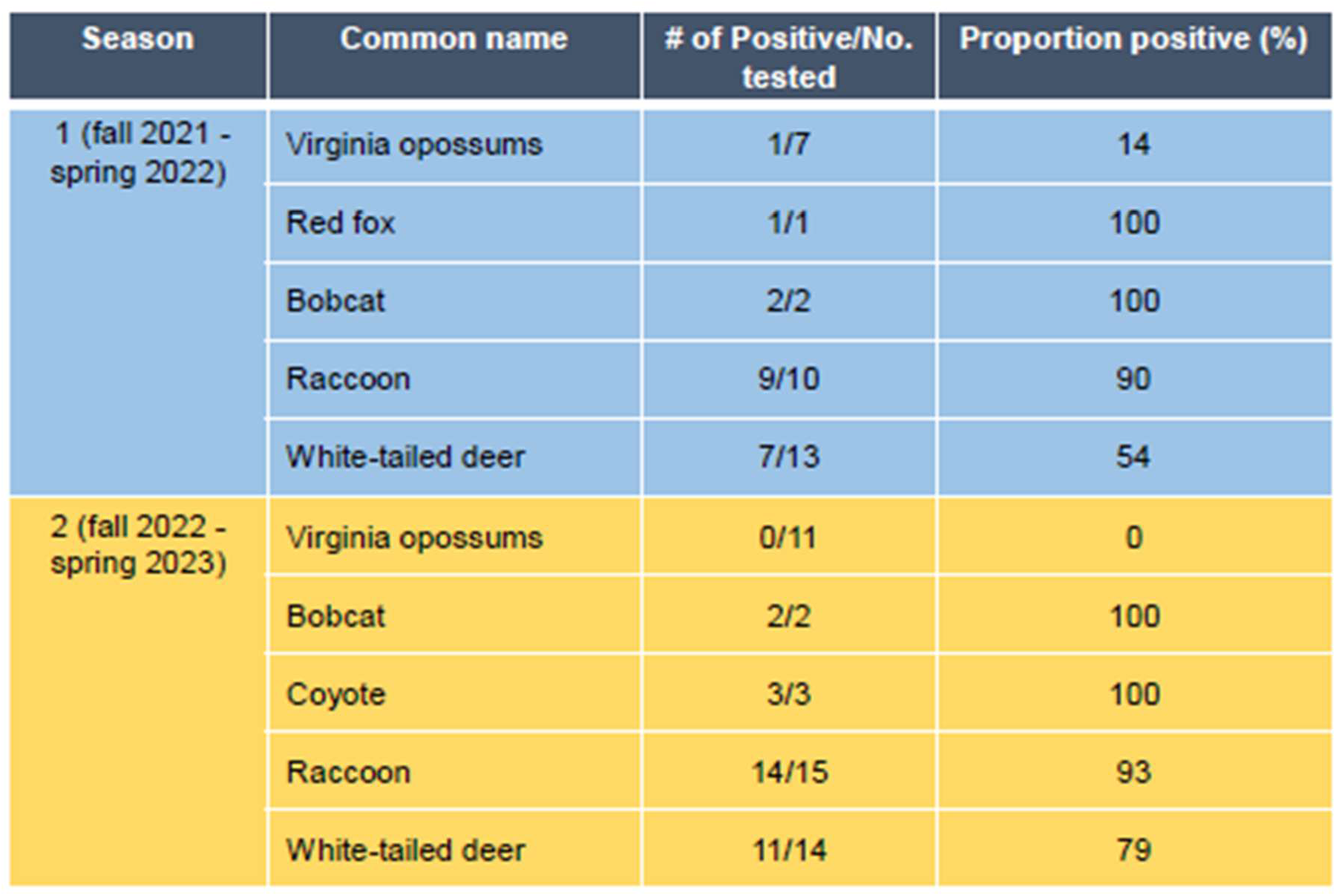
Number and frequency of BRBV-neutralizing antibody-positive animals in TRC between Fall 2021 and Spring 2023.

| Season | Common name | # of Positive/No. tested | Proportion positive (%) |
| --- | --- | --- | --- |
| 1 (fall 2021 - spring 2022) | Virginia opossums | 1/7 | 14 |
|  | Red fox | 1/1 | 100 |
|  | Bobcat | 2/2 | 100 |
|  | Raccoon | 9/10 | 90 |
|  | White-tailed deer | 7/13 | 54 |
| 2 (fall 2022 - spring 2023) | Virginia opossums | 0/11 | 0 |
|  | Bobcat | 2/2 | 100 |
|  | Coyote | 3/3 | 100 |
|  | Raccoon | 14/15 | 93 |
|  | White-tailed deer | 11/14 | 79 |

**Figure 1:**
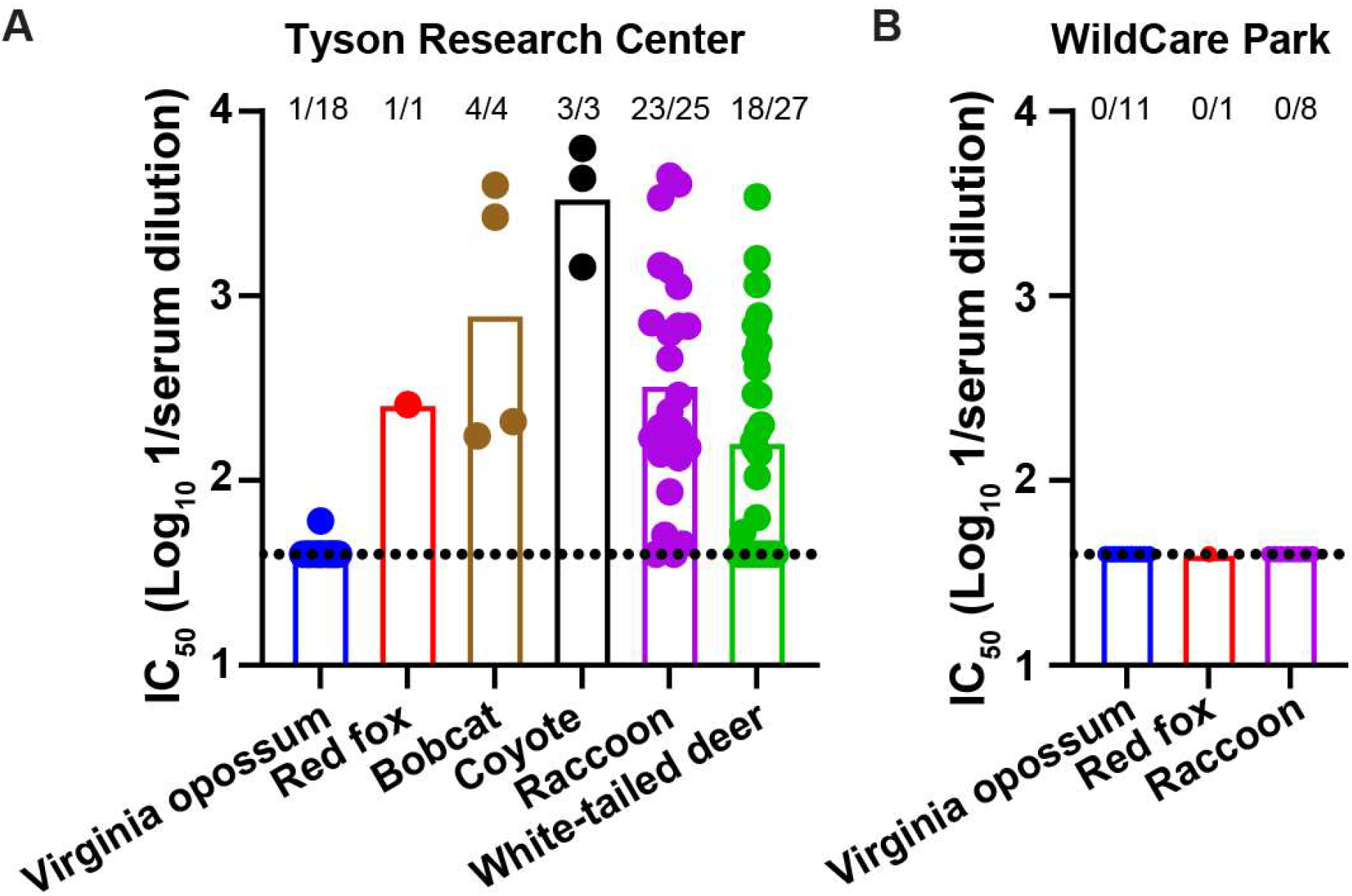
Detection of Bourbon virus neutralizing antibodies in diverse wildlife species. Serum samples collected from Virginia opossums (n = 29), red foxes (n = 2), bobcats (n = 4), coyotes (n = 3), raccoons (n = 33), and white-tailed deer (n = 27) between Fall 2021 and Spring 2023 in Tyson Research Center (A) and WildCare Park (B) were tested for the presence of BRBV neutralizing antibodies by focus reduction neutralization assay. The serum dilution inhibiting 50% of the virus (IC_50_) is plotted for each individual and the number of BRBV seropositive individuals are indicated above the bars. The dotted line is the limit of detection for the assay.

### Virus detection in sera from wildlife

Sera collected were also tested for the presence of BRBV RNA by RT-qPCR using primers and probe sets targeting the nucleoprotein (NP) and polymerase basic protein 1 (PB1) of BRBV. Out of the 98 samples tested, one coyote serum sample was positive for both NP and PB1 with Ct values of 33.1 and 31.5 respectively. To validate our findings, we performed whole gene-segment RT-PCR using segment-specific primers targeting the 5’ and 3’ untranslated regions (UTR) of all six segments. Full-length amplicons were obtained for BRBV PB1, glycoprotein (GP), and NP. Sanger sequencing of these amplicons confirmed they were BRBV gene segments. More importantly, four synonymous nucleotide polymorphisms at position 954, 963, 969, and 972 were identified in GP that have not been found in any other BRBV sequence or isolate to date. No changes were detected in the NP and PB1 gene amplified from the coyote serum sample.

The BRBV-positive female coyote was captured on January 23, 2023, in TRC. Physical examination at the time of capture was considered within normal limits based on a body weight of 14 kg; and hydration status, appearance of fur coat, body condition score, and absence of clinical signs of disease in all organ systems explored during the exam. Additionally, no external lesions were observed in the individual and mammary glands and teats were undeveloped, so we concluded this female was nulliparous. White blood cell differential was non-remarkable with segmented neutrophils (77%), lymphocytes (14%), eosinophils (6%), monocytes (3%), and basophils (0%) [30].

### Field Collected Ticks

Using the same trapping methods, we collected 2,323 ticks at TRC and 31 ticks at WCP (Table 3). All TRC traps yielded *A. americanum* ticks in both sampling rounds (98.4%), and nymphs (77%) and adults (21%) were the predominant life stages captured. Of these adult ticks, 38% were males and the rest were females (62%). The remaining ticks were *Dermacentor variabilis* (1.3%) and *Ixodes scapularis* (0.3%). The 31 ticks at WCP included 26 larval *A. americanum* from only one trap during one sampling round, and 5 adult *D. variabilis*.

**Table 3.** Information on ticks collected in the study on BRBV in Missouri wildlife. **Table 3:** Number of collected ticks in four different sites at Tyson Research Center (TRC) and WildCare Park (WCP). The data are separated by tick species (AA = *Amblyomma americanum*, DV = *Dermacentor variabilis*, IS = *Ixodes scapularis*), sex, and life stage (AA only).

| Location | Tick species |  |  |  |  |  |  |
| --- | --- | --- | --- | --- | --- | --- | --- |
|  | AA |  |  |  |  | DV | IS |
|  | Total | Male | Female | Nymph | Larva | Total | Total |
| **TRC** |  |  |  |  |  |  |  |
| Site 1 | 1328 | 87 | 119 | 1122 | 0 | 4 | 1 |
| Site 2 | 385 | 50 | 85 | 250 | 0 | 12 | 6 |
| Site 3 | 315 | 29 | 61 | 225 | 0 | 6 | 0 |
| Site 4 | 255 | 12 | 24 | 219 | 0 | 3 | 1 |
| Total | 2283 | 178 | 289 | 1816 | 0 | 25 | 8 |
| **WCP** |  |  |  |  |  |  |  |
| Site 1 | 0 | 0 | 0 | 0 | 0 | 1 | 0 |
| Site 2 | 0 | 0 | 0 | 0 | 0 | 3 | 0 |
| Site 3 | 26 | 0 | 0 | 0 | 26 | 1 | 0 |
| Site 4 | 0 | 0 | 0 | 0 | 0 | 0 | 0 |
| Total | 26 | 0 | 0 | 0 | 26 | 5 | 0 |

**Table 4:**
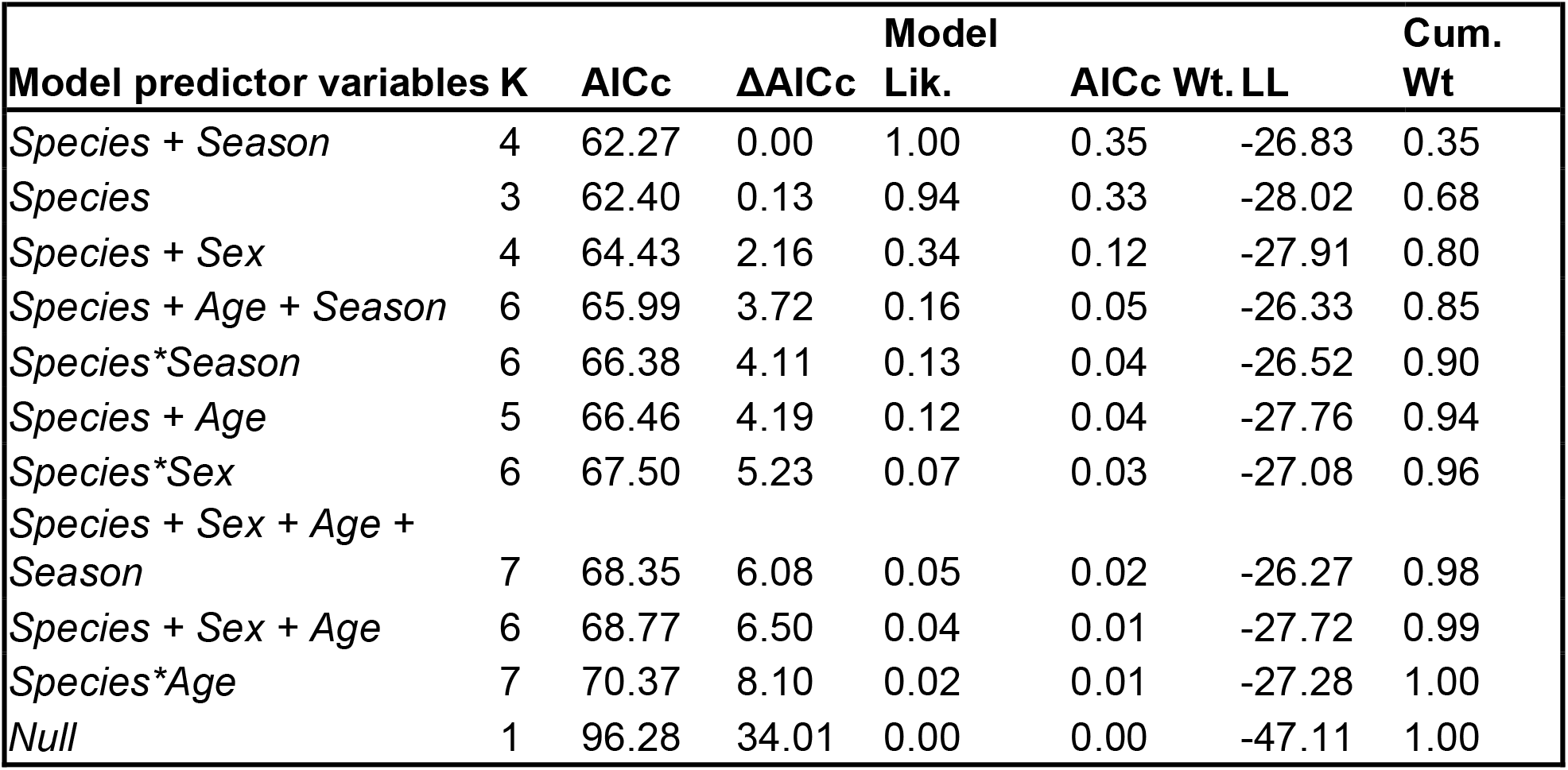
AICc Model Selection Results in the study of BRBV in Missouri wildlife. **Table 4:** Predicted odds of an individual white-tailed deer, raccoon, or Virginia opossum being seropositive for BRBV at Tyson Research Center using a set of 11 logistic regression models.

### Predictors of seropositivity

Among white-tailed deer, opossums, and raccoons sampled from TRC, the most influential variables to describe the likelihood of an individual being positive for BRBV antibodies were species, season, and sex. The top three models, determined by a threshold of roughly ΔAICc = 2.0 from among the candidate sets, carried 80% of the cumulative model weight (Table 4). Each of the three models included species as a predictor variable, and the univariate species model outperformed the null model by ΔAICc > 30. While almost no opossums were seropositive (mean model-predicted probability of seropositivity = 5.6%, [95%CI: 0.78%-30.7%]), raccoons (mean predicted probability = 92.0%, [73.1% - 98.0%]) were more likely to be seropositive than deer (mean predicted probability = 66.7%, [47.3% - 81.7%]). Top models also estimated a mean ∼5% increase in the probability of seropositivity across all individuals between seasons 1 and 2, suggesting increasing exposure rates to BRBV at TRC over time. Male animals were on average less likely to be seropositive than females, but there was a large degree of uncertainty around the odds ratio estimate for male seropositivity relative to females (0.714 [95%CI: 0.17-2.91]). We note that models containing season and sex as predictors performed similarly to the species-only model (Table 4), indicating that capture season and host sex had limited predictive power relative to species for this small dataset. Model selection did not identify host age as an important predictor of BRBV seropositivity.

## DISCUSSION

We report for the first time BRBV seropositive bobcats, coyotes, and red foxes, adding to the list of species previously reported to be seropositive for BRBV. Furthermore, we detected BRBV RNA in the serum of a coyote, indicating that the animal may have been viremic for BRBV at the time of capture. This is the first report of BRBV genetic material isolated from any wildlife species. As in prior studies [6, 18, 19], raccoons and white-tailed deer exhibited high BRBV seropositivity rates, while only 1/18 Virginia opossums were positive for BRBV-specific antibodies in our cohort of animals captured at TRC, St. Louis, MO. We identified an increasing exposure risk of BRBV in raccoons and white-tailed deer across two years at TRC. In a comparison of two field sites within the St. Louis, MO area, we observed wildlife seropositivity only in the location with greater lone star tick density, suggesting localized wildlife exposure to BRBV. Collectively our results contribute to a broader understanding of BRBV ecology and expand the pool of potential wildlife species involved in BRBV maintenance and transmission.

Previous serosurveillance studies from Missouri, North Carolina, and New York detected BRBV exposure in white-tailed deer, with seroprevalence rates from 38-86% detected by serum-neutralizing antibodies [6, 18, 19]. Similarly, sera collected at TRC, St. Louis, MO, also showed high seroprevalence rates among deer in the two capture seasons (54% and 79%). White-tailed deer have long been important sentinels for tick populations and tick-borne disease surveillance. Our results underscore the value of continued deer monitoring for BRBV surveillance. Partnering with hunters, conservation agencies, and deer serum repositories may allow a much broader assessment of BRBV distribution. Jackson *et al* 2019 observed a 50% seroprevalence rate in raccoon sera collected from northwestern MO [18]. Here, raccoons at TRC had the highest prevalence of BRBV-neutralizing antibodies (90 and 93%), which was greater than the prevalence rate observed for white-tailed deer. In contrast, all raccoons captured at WCP were negative for BRBV exposure. These findings demonstrate that white-tailed deer and raccoons have a high seroprevalence and could potentially serve as local sentinels for BRBV presence.

For the 1 red fox, 4 bobcats, and 3 coyotes sampled at TRC, all were seropositive for BRBV. The sample sizes from these species were too small to analyze statistically, but their apparent high seropositivity rates and high antibody titers are notable. Relative to white-tailed deer and raccoons, these species’ lower population densities and capture success rates make them less useful as potential sentinels; however, much remains unknown about the transmission ecology of BRBV. We detected BRBV RNA by RT-qPCR in a sample from an adult female coyote that was captured on January 24, 2023, which is outside of the typical lone star tick activity season in Missouri. This finding raises questions about the possible timing and duration of the coyote’s BRBV infection, its potential as a reservoir, or amplifying, host, and whether other tick species might be involved in BRBV transmission.

Despite the high seropositivity rates for multiple species at TRC, we found only one Virginia opossum positive for BRBV-neutralizing antibodies. Similarly, Jackson *et al* 2019 did not find any BRBV seropositive individuals out of a sample of 28 opossums in Missouri [18]. The reason for the lack of BRBV antibodies in Virginia opossums is not known, but one possibility is that opossums feed fewer ticks relative to other host species, as reported previously [17, 31]. However, contrary to this evidence, a few more studies have identified Virginia opossums as suitable hosts for several tick species, including lone star ticks [32, 33]. Another possibility is that opossums are less susceptible to BRBV infection due to more effective innate immune responses, lower body temperatures [34], or the lack of a critical host factor required for virus replication. Collectively, these findings suggest that the BRBV infection rates in Virginia opossums are low. However, the mechanism behind these findings warrants further investigation.

Compared to TRC with its abundant lone star tick population, we found very few lone star ticks in WCP in 2023. Although the serum sample size from WCP was limited (n = 20), the lack of BRBV positivity in all raccoons is likely related to the small lone star tick population and we consider that BRBV may not be present at this site. Habitats at WCP differ from those at TRC, and it may be important to conduct continuous monitoring of hosts, tick density, and BRBV seroprevalence in WCP over time to understand how each factor influences BRBV ecology. BRBV is present in ticks at TRC [11] and hosts have high exposure rates, which makes TRC an ideal site to continue monitoring BRBV seroprevalence and study BRBV transmission and viremia through continued sampling, collection of ticks parasitizing susceptible host species during peak lone star activity periods, and broadening the diversity of species sampled to further our understanding of BRBV ecology, maintenance and transmission.

### Limitations to the study

There are several limitations to the study. First, we did not sample the same number of animals and species at WCP, limiting our ability to model predictors of seropositivity. Second, we were unable to sequence the full genome of BRBV from the coyote serum sample. Third, we did not perform RT-qPCR analysis on ticks collected from the captured wildlife species. Finally, the absence of neutralizing antibodies does not rule out that these animals have never been exposed to BRBV. Waning immunity or low-level infection can cause neutralizing antibody titers to fall below the limit of detection.

Collectively, through our study, we identified three additional mammalian hosts susceptible to infection with BRBV and detected the first BRBV viremia in a coyote. Tick density, wildlife species, and year of sampling are important risk factors for BRBV exposure, and raccoons and white-tailed deer are excellent sentinels for BRBV serosurveillance. Continued surveillance of BRBV, ticks, and wildlife species will broaden our understanding of BRBV ecology, spread, and distribution.

## ACKNOWLEDGEMENTS

We thank Tyson Research Center and the Saint Louis Zoo WildCare Institute for funding and Susan Flowers for program support. Undergraduate students Althea Bartz Willis, Yoshihiro Yajima, and Chenxi Zhang assisted with tick collections and identification as part of the Tyson Undergraduate Fellowship Program. We also thank the following hunters who provided samples from white-tailed deer: Pete Jamerson, Tim Derton, Mike Parker, Doug Parker, Chris Parker, Mike Mueller, Darrell, Beau, Patrick Maschmeyer, Steve Maschmeyer, Cameron Miles, Carl Vanderloo, and Dan Dudley. This work was supported by U01-AI151810 (ACMB and DW).

